# MeShClust v3.0: High-quality clustering of DNA sequences using the mean shift algorithm and alignment-free identity scores

**DOI:** 10.1101/2022.01.15.476464

**Authors:** Hani Z. Girgis

## Abstract

**Background:** Tools for accurately clustering biological sequences are among the most important tools in computational biology. Two pioneering tools for clustering sequences are *CD-HIT* and *UCLUST*, both of which are fast and consume reasonable amounts of memory; however, there is a big room for improvement in terms of cluster quality. Motivated by this opportunity for improving cluster quality, we applied the mean shift algorithm in *MeShClust v1.0*. The mean shift algorithm is an instance of unsupervised learning. Its strong theoretical foundation guarantees the convergence to the true cluster centers. Our implementation of the mean shift algorithm in *MeShClust v1.0* was a step forward; however, it was not the original algorithm. In this work, we make progress toward applying the original algorithm while utilizing alignment-free identity scores in a new tool: *MeShClust v3.0*.

**Results:** We evaluated *CD-HIT, MeShClust v1.0, MeShClust v3.0*, and *UCLUST* on 22 synthetic sets and five real sets. These data sets were designed or selected for testing the tools in terms of scalability and different similarity levels among sequences comprising clusters. On the synthetic data sets, *MeShClust v3.0* outperformed the related tools on all sets in terms of cluster quality. On two real data sets obtained from human microbiome and maize transposons, *MeShClust v3.0* outperformed the related tools by wide margins, achieving 55%—300% improvement in cluster quality. On another set that includes degenerate viral sequences, *MeShClust v3.0* came third. On two bacterial sets, *MeShClust v3.0* was the only applicable tool because of the long sequences in these sets. *MeShClust v3.0* requires more time and memory than the related tools; almost all personal computers at the time of this writing can accommodate such requirements. *MeShClust v3.0* can estimate an important parameter that controls cluster membership with high accuracy.

**Conclusions:** These results demonstrate the high quality of clusters produced by *MeShClust v3.0* and its ability to apply the mean shift algorithm to large data sets and long sequences. Because clustering tools are utilized in many studies, providing high-quality clusters will help with deriving accurate biological knowledge.

## Background

Clustering DNA sequences has broad applications in molecular biology. Clustering tools have been applied to grouping transposable elements [1–4], open reading frames [5, 6], and expressed sequence tags [7–10]. Cluster analysis has been utilized as a “complement” to phylogenetic analysis [11]. Clustering tools have been used in identifying sub-types in a viral population [12], identifying “non-reference representative sequences” needed for constructing pangenome [13], decomposing genomes [14], and “assigning individuals” to operational taxonomic units using DNA barcodes [15].

The two most widely used clustering tools are *CD-HIT* [16] and *UCLUST* [17]. These pioneering tools have been utilized in thousands of studies. *CD-HIT* and *UCLUST* depend on greedy algorithms for clustering sequences and on the Needleman-Wunsch global alignment algorithm [18, 19] for evaluating sequence similarity. While studying these two tools, we identified two main limitations. Because of the greedy nature of these tools, they are not guaranteed to find the optimal clusters, i.e., the true clusters. The clusters produced by *CD-HIT* and *UCLUST* are likely fragments of the true clusters. Because of the slow nature of the global alignment algorithm, the two tools cannot be applied to clustering very long sequences such as bacterial genomes.

Motivated to overcome the first limitation, we proposed *MeShClust v1.0* [20], which is based on the mean shift algorithm [21]. The ability of the mean shift algorithm to find the true cluster centers is proven theoretically. There are more than 15,000 applications of the mean shift algorithm in the computer vision field [22–24]. Our bioinformatics field is slowly taking advantage of this powerful algorithm [20, 25–28]. The first adaptation of the mean shift algorithm to clustering DNA sequences resulted in clusters of better quality than those produced by *CD-HIT* and *UCLUST* [20]. This adaptation is not the orthodox implementation of the mean shift algorithm. What prevented us from applying the orthodox mean shift algorithm in *MeShClust v1.0* is its initialization step that requires evaluating similarity between every two sequences in a data set. Such initialization step would take an impractical long time on large data sets. We are convinced that the orthodox mean shift algorithm should result in high-quality clusters. To this end, we propose a new adaptation of the mean shift algorithm that is closer to the orthodox algorithm than our earlier adaptation.

*MeShClust v1.0* overcame the first limitation of *CD-HIT* and *UCLUST*; however, it cannot be applied to very long sequences because it is assisted by a global alignment algorithm. *CD-HIT*, *MeShClust v1.0*, and *UCLUST*, utilize identity scores as the sequence similarity measure. We define an identity score of two sequences as the percentage or ratio of the number of identical nucleotides in two optimally aligned sequences to the length of the alignment. The alignment length can be longer than the length of any of the two sequences because an alignment may include gaps representing insertions and deletions. Traditionally, identity scores are produced by alignment algorithms. Recently, we have devised an efficient machine-learning-based alternative to global alignment algorithms in a tool we call *Identity* [29]. *Identity* can produce pairwise identity scores efficiently and without performing any costly alignments. Taking advantage of *Identity, MeShClust v3.0* can cluster long sequences such as those of bacterial genomes, overcoming the second limitation.

In sum, the two main contributions of *MeShClust v3.0* are: (i) high quality clusters and (ii) the ability to cluster long sequences using alignment-free identity scores.

## Implementation

Here, we explain how the mean shift algorithm works. Then we propose a new out-of-core learning strategy in order to scale the algorithm to large data sets.

The original algorithm and the scaled-up version require a method for calculating the similarity between two sequences; in this work, we utilize identity scores. We could use the Needleman-Wunsch global alignment algorithm for calculating identity scores; however, this algorithm is quadratic and will not scale to a large number of sequences or very long sequences. For this reason, we utilize a machine-learning-based tool called *Identity* for predicting identity scores much faster than the Needleman-Wunsch global alignment algorithm (linear vs. quadratic).

### Calculating alignment-free identity scores efficiently

*Identity* is designed to work on large sequence sets and long sequences, e.g., bacterial genomes. *Identity* is an instance of self-supervised learning. The tool selects sequences from the input set to generate its own labeled training data. Each sequence of the selected input sequences is mutated to produce multiple similar copies with known identity scores when compared to the original sequence. The idea is that *Identity* keeps track of the mutations (introduced to a copy of the original sequence), each of which could change the alignment length and the number of matched nucleotides in a unique way if the original sequence and the mutated version were to be aligned. For example, introducing an insertion increases the length of the alignment by the size of the insertion and has no effect on the number of matched nucleotides. This way, the identity score can be calculated *without applying an alignment algorithm* by dividing the number of matches by the length of the alignment; both numbers are calculated according to the mutations introduced to the original sequence.

At this point, *Identity* has a set of sequence pairs with their identity scores; it is ready to train itself. The training process is automatic and involves the following steps:

- Convert each sequence to a k-mer histogram (*k* = ⌈log_4_ average length of training sequences⌉ — 1).
- Calculate a set of single statistics, e.g., Manhattan distance, on each pair of histograms representing an original-mutated sequence pair; take the square of each statistic to produce squared statistics; multiple every unique pair of statistics (single or squared) to produce paired statistics.
- Select a subset of these statistics (single, squared, and paired) using the best-first algorithm [30].
- Train a general linear model on the selected statistics, resulting in a weight for each selected statistic.

Now, *Identity* is ready to predict the identity score of any sequence pair in an input data set. Two sequences are converted to two k-mer histograms, on both of which the selected statistics are calculated. The identity score is calculated as the weighted sum of the statistics.

*Identity* has an option to skip a sequence pair if it is impossible to produce a desired minimum identity score. A sequence pair is skipped if (i) the length ratio (length of short sequence ÷ length of long sequence) is less than the desired identity score or (ii) their monomer potential (Equation 3) is less than the desired threshold score.

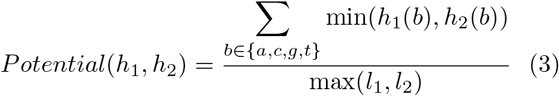

Here, *h*_1_ and *h*_2_ are monomer histograms of two sequences, and *l*_1_ and *l*_2_ are the lengths of the two sequences. This equation is similar to the one applied in the MUSCLE program [31]. We apply it in *Identity* to monomer histograms, whereas it is applied to k-mer histograms in MUSCLE.

Note that the user is not required to provide labeled training data; *Identity* generates its own training data from the input sequences. Further, the training process is repeated on each input sequence set; the process is automatic and does not require any involvement from the user. Next, the mean shift clustering algorithm utilizes *Identity* for calculating pairwise identity scores.

#### Algorithm 1 Sequence clustering using the mean shift algorithm

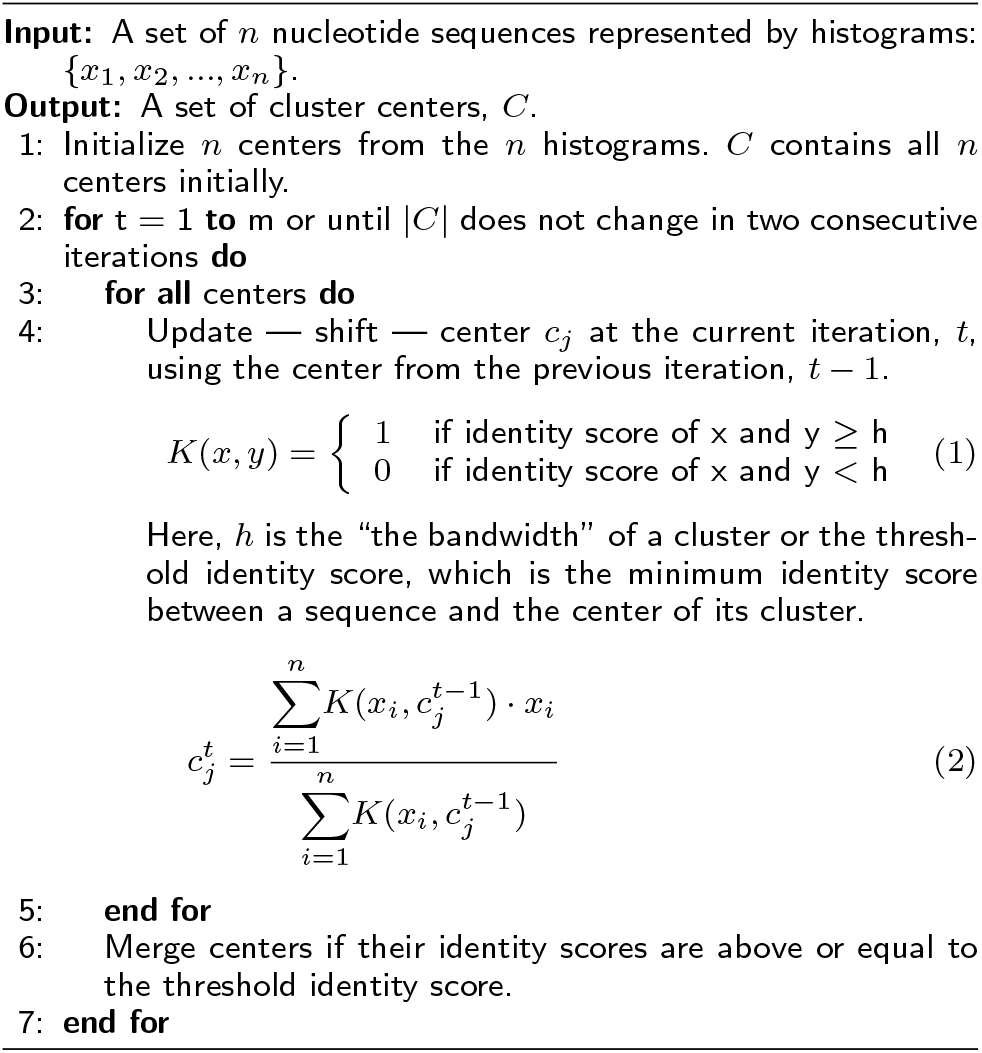

### Clustering few thousands of sequences

The original mean shift algorithm — without any modifications — can cluster 25K–45K sequences in reasonable time using memory available on personal computers. The algorithm is iterative and consists of four steps. The first step is executed only once, but the second, third, and fourth steps are executed many times until the algorithm converges. Algorithm 1 outlines the mean-shift algorithm as it is applied to nucleotide sequences. Input sequences are converted to k-mer histograms (k is determined the same way as in *Identity*), on which the algorithm performs the following steps:

- Step 1: Initialize. Every k-mer histogram is treated as a center of a one-member cluster.
- Step 2: Shift. A new mean for each cluster is computed as the average of sequences (k-mer histograms) that have identity scores with the center of the cluster above or equal to a threshold score (provided by the user or estimated automatically).
- Step 3: Merge. Centers whose identity scores are greater than or equal the threshold identity score are merged. We keep the center that represents the mean of the largest number of histograms.
- Step 4: Represent. The mean of each cluster is represented, i.e., replaced, by the sequence (k-mer histogram) with the highest identity score.

Steps 2–4 are repeated until the algorithm converges. Note that Step 3 and Step 4 are reversed when clustering degenerate sequences (≤ 70% identity score); this way, the merging step is performed on k-mer histograms of real sequences not on the means of degenerate sequences. Convergence occurs when the number of iterations is reached or the number of clusters does not change in two consecutive iterations.

Once the algorithm converges, we have centers — but no clusters yet. Input sequences are re-read. Each sequence is assigned to the center that has the highest identity score with it.

### Clustering hundreds of thousands of sequences

To run the original mean shift algorithm on a large data set would take impractically long time because of the initialization step. In the original algorithm, each sequence is treated as the center of a one-member cluster in the initialization step. Then the identity score of every two centers is calculated. This step is quadratic with respect to the number of sequences in a data set — very slow on very large data sets. To make the issue clear, consider a data set consisting of 1,000,000 sequences. The initialization step requires 499,999,500,000 sequence comparisons. Calculating this huge number of identity scores — even with an efficient tool such as *Identity* — takes approximately one day. Keep in mind that this number of sequence comparisons is required in the first step only, and more sequence comparisons will be performed in subsequent steps.

To circumvent this limitation, we propose an out-of-core learning adaptation to the original mean shift algorithm. In out-of-core learning, the learning algorithm (the mean shift clustering algorithm here) is trained on batch by batch instead of the entire data set [32]. The large-scale mean shift algorithm requires multiple passes through the data.

To allow the algorithm to work on batch by batch, Step 2 (the mean-shift step) needs to be modified. The algorithm keeps track of how many sequences (k-mer histograms) contributed to each center. In the modified Step 2, the new mean is calculated as the weighted average of the old mean and the new sequences. For example, suppose that 80 sequences contributed to a center after the algorithm had run on some batches. While processing the next batch, 20 sequences were similar to the center (their identity scores with the center were above or equal to the threshold score). The new mean is the weighted average of the old mean (weight: 80/100 = 0.80) and each of the new 20 sequences (weight: 1/100 = 0.01). Once the new mean is computed, the number of sequences contributing to the mean is updated to 100 (80 + 20).

We outline the first data pass of the out-of-core mean shift algorithm in Figure 1. To begin, a batch of the input sequences is read. Once the first batch is read, the four steps of the original mean shift algorithm are executed until convergence. Sequences that do not belong to any cluster of size greater than one — we call them singles — are kept in a reservoir. After that, a new batch is read. The algorithm is initialized using the centers found in the previous batch(es), i.e., no new centers are introduced. Steps 2–4 are executed on the new batch one time. Next, singles found in the new batch are added to the reservoir. After processing a batch, *MeShClust v3.0* checks to see if enough (more than the batch size) sequences have been accumulated in the reservoir. If yes, sequences in the reservoir are shuffled; a new instance of the mean shift algorithm is executed on a batch obtained from the reservoir — not from the input file. This instance is initialized according to the original algorithm, i.e., every sequence represents the center of a one-member cluster. It is run until convergence. Centers found in this batch are added to the main mean shift instance. Some of the centers found in the reservoir batch may be merged with some of the centers found so far; others are new centers. To fill the reservoir faster, the number of sequences to read increases adaptively by a factor equal to the batch size (25,000 is the default) divided by the number of singles found in the previous sequences. The increase is limited by a maximum of four times, i.e., the maximum number of sequences that can be read when using the default batch size is 100,000 sequences. These steps are repeated until all sequences in the input file are processed and the reservoir is empty.

**Figure 1:**
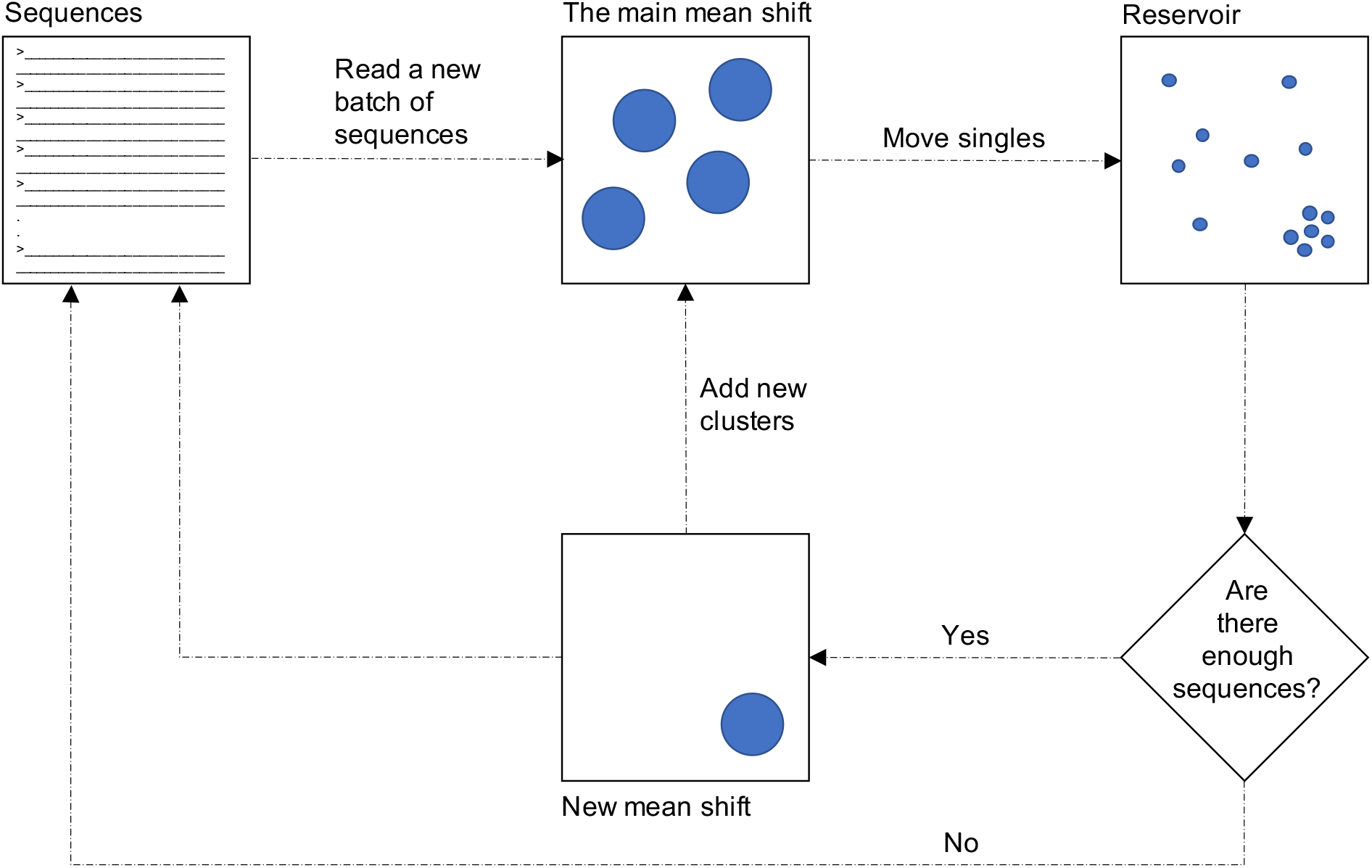
Overview of the first data pass. *MeShClust v3.0* is based on the mean shift algorithm, which is an instance of unsupervised learning. The scaled-up *MeShClust v3.0* is also an instance of out-of-core learning [32], in which the learning algorithm is trained on separate batches of the training data consecutively. The algorithm requires multiple passes through the input data. In the first data pass, the tool reads a batch of input sequences. Then the mean shift algorithm (all of the four steps) is run on the batch until convergence. Sequences that cannot be assigned to any center are kept in the reservoir. Next, a new batch is read. The main mean shift is run on this batch but without the initialization step and for one iteration only, i.e., already found centers are shifted and merged on the new batch and no new centers are discovered. Sequences that cannot be assigned to any of the centers are added to the reservoir. When the reservoir has enough sequences (more than the default batch size), sequences in it are shuffled and a batch of them is clustered using an independent instance of the mean shift algorithm. This instance is run until convergence. The resulting centers (if any) are merged with the centers accumulated by the main mean shift. This procedure is repeated until all sequences are read and the reservoir is empty. In subsequent passes, the algorithm rereads input sequences batch by batch. The main mean shift algorithm is run for one iteration on each batch. If the number of clusters does not change during a pass, the algorithm converges. In the final data pass, all sequences are reread batch by batch, and each sequence is assigned to the cluster with the closest center to it.

In subsequent data passes, input sequences are reread batch by batch. Steps 2–4 are executed once on each batch. Singles are *not* added to the reservoir. If the number of centers does not change in a data pass, the algorithm converges.

In the final data pass, input sequences are re-read batch by batch. Each sequence is assigned to the cluster with the closest center to it.

Overall, the first data pass is for discovering centers, the subsequent passes are for fine-tuning them, and the final pass is for forming clusters.

This adaptation was motivated by the need to reduce the number of sequence comparisons at the initialization step. The number of sequence comparison required for the initialization step on a 25K-sized batch is 312,487,500. *Identity* can calculate these scores in a matter of seconds or minutes — not a day as it is the case if the original algorithm was to be applied. The rational is that a sequence that belong to one of the already found clusters does not need to form its own cluster because this cluster would merge with the already-found cluster. Further, in a data set that has a smaller number of clusters than the batch size, most of the clusters could be found by running the mean shift on the first batch. In case some of the clusters are missed, we accumulate singles in the reservoir. Then, sequences in the reservoir are clustered independently to find any clusters that could have been missed.

### Estimating the threshold score

One advantage of *MeShClust v3.0* is that the user may choose not to provide a threshold score. In this case, the tool determines the threshold score from input data automatically. The idea is that for a sequence to belong to a cluster, its identity score with the center must be greater than or equal to the threshold score. If we assume that any sequence may represent the center of a cluster and the smallest cluster consists of five sequences, then the threshold score is approximately equal to the maximum identity score between the center and any of the other four members.

To determine the maximum center-member identity score, *MeShClust v3.0* reads 10,000 sequences. It calculates all-versus-all identity scores on these sequences using *Identity*. For each sequence, the highest four identity scores are collected.

If the size of the input data set is smaller than 10,000, the threshold score is estimated to be the mean minus three times the standard deviation. Assuming that these scores are normally distributed, this threshold score should cover 99.9% of the data. If the size of the input data set is greater than 10,000, the collected scores are clustered into two groups. One group should represent scores of sequences that belong to clusters. The other group should represent sequences that do not belong to clusters; this case is possible if the data include noise or the batch analyzed is smaller than the entire input. *MeShClust v3.0* utilizes the k-means clustering algorithm with a k value of 2. Assuming that the percentage of noisy scores is small, the cluster with the smallest number of points is removed. The estimated threshold, *t*, is determined by Equation 4.

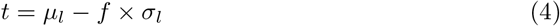

Here, *μ_l_* and *σ_l_* are the mean and the standard deviation of the large cluster, and *f* is determined according to the ratio of the batch size to the input size.

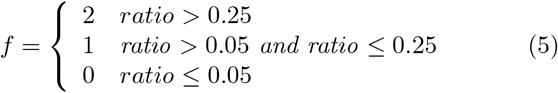

This estimation procedure is repeated three times (if the input data are large enough) using a new batch each time; the median threshold is selected.

### Related tools

We compared the performance of our tool to those of *CD-HIT* [16], *MeShClust v1.0* [20], and *UCLUST* [17]. *CD-HIT* was run with -M with a value of 0, -G with a value of 1, -d with a value of 0, -T with a value of 16, and -c with a value of the desired threshold score. If the threshold identity score is between 60% and 50%, the -t parameter was used with a value of 3. When the threshold identity score is less than 50%, the -n parameter was used with a value of 3. *MeShClust v1.0* was run with the “threads” parameter with a value of 16. For *UCLUST*, the usearch program was executed with -cluster_fast, -id with a value of the desired threshold score, -sort with a value of “length”, and -threads with a value of 16. *MeShClust v3.0* was run with -t with a value of the desired threshold score. In the experiments where *MeShClust v3.0* estimated the threshold score itself, it was run without any parameters. *MeShClust v3.0* determines the number of threads automatically as the number of hyper-threads available on a computer, i.e., *MeShClust v3.0* utilized 16 threads.

### Evaluation computer

All experiments were conducted on a computer with 8 cores/16 hyper-threads (16 MB Cache and 3.6 Ghz), Nvidia Quadro RTX 4000 graphic card, 64 GB of RAM, 1-TB solid state disk, two 2-TB hard disk drives, and Ubuntu 18.04 as the operating system.

### Data

To evaluate *MeShClust v3.0* and the related tools, we collected real sequences and generated synthetic ones.

#### Real Data

Five real data sets were utilized in this work. These data sets represent bacterial genomes, viral genomes, the 16S rRNA gene of the human microbiome [33], and Long Terminal Repeats (LTRs) of maize retrotransposons. The viral set was obtained from viruSITE [34]. All bacterial genomes were downloaded from the NCBI. Table 1 shows statistics on these data sets.

**Table 1:** Statistics of the real data sets. The 14-bacterial-species data set includes 14 clusters. The viral set includes 9 clusters. Cluster counts in the bacterial, maize LTRs, and human microbiome sets are unknown.

| Data set | Sequence count | Total length | Maximum length | Minimum length | Mean length | Median length |
| --- | --- | --- | --- | --- | --- | --- |
| 14 bacterial species | 1,328 | 4,256,374,969 | 9,270,175 | 801,203 | 3,205,102 | 2,874,351 |
| Bacterial | 10,562 | 38,577,794,947 | 16,040,666 | 112,031 | 3,652,509 | 3,647,501 |
| LTRs | 253,224 | 346,337,915 | 5,999 | 100 | 1,368 | 1,187 |
| Microbiome | 1,071,335 | 269,374,512 | 372 | 171 | 251 | 256 |
| Viral | 96 | 635,979 | 13,246 | 2,605 | 6,625 | 7,458 |

#### Synthetic Data

One of the advantages of using synthetic data sets is that we know the true clusters and their true centers. We generated 22 *training* data sets, on which we trained and optimized *MeShClust v3.0*, and additional 22 *testing* data sets, on which we evaluated our tool and the related ones. Table 2 shows the statistics of the training data sets. The testing data sets have very similar statistics to the training ones (data are not shown). To generate a data set, a number of random sequences — templates — are synthesized. These templates may represent the true cluster centers. The length of each template is chosen at random between a minimum length and a maximum length. To reduce the chance of generating intersecting clusters, identity scores among all templates in the same data set are at most 10% less than the provided threshold score that determines cluster membership. A random number (between a minimum and a maximum) of mutated copies are generated from each template by introducing single-point mutations and block mutations to a copy of a template. These mutated copies are generated using the module that generates training data for *Identity* [29]. All mutated copies generated from the same template have identity scores with the template greater than or equal to the provided threshold score. Each of the small data sets (Short-X, Medium-X, and Long-X) includes 100 clusters and less than 25,000 sequences. Each of the large data sets (Numerous-X) includes 5,000 clusters and about one million sequences.

**Table 2:** Statistics of the synthetic training data sets. To construct a synthetic data set, a specific number of template random sequences are synthesized. The length of a template is chosen at random between minimum and maximum lengths. A random number (between minimum and maximum numbers) of mutated copies are generated from each template. All clusters in the same data set have the same minimum identity score. For example, members comprising the clusters of the Short-97 data set are 97.00–99.99% identical to the templates, from which these members were generated. Identity scores among templates in the same data set are at most 10% less than the provided minimum identity score.

| Data set | Template avg. length (bp) | Template min. length (bp) | Template max. length (bp) | Cluster avg. size | Cluster min. size | Cluster max. size | Cluster count | Sequence count |
| --- | --- | --- | --- | --- | --- | --- | --- | --- |
| Short-97 | 288 | 202 | 396 | 202 | 12 | 400 | 100 | 20,195 |
| Short-95 | 307 | 200 | 400 | 177 | 5 | 400 | 100 | 17,734 |
| Short-90 | 298 | 204 | 399 | 199 | 9 | 400 | 100 | 19,877 |
| Short-80 | 302 | 200 | 400 | 204 | 6 | 392 | 100 | 20,423 |
| Short-70 | 299 | 205 | 400 | 202 | 7 | 395 | 100 | 20,230 |
| Short-60 | 304 | 200 | 399 | 195 | 9 | 395 | 100 | 19,539 |
| Medium-97 | 1,394 | 752 | 1,998 | 192 | 13 | 390 | 100 | 19,215 |
| Medium-95 | 1,358 | 750 | 1,968 | 203 | 7 | 396 | 100 | 20,315 |
| Medium-90 | 1,405 | 759 | 1,977 | 194 | 5 | 400 | 100 | 19,393 |
| Medium-80 | 1,434 | 760 | 2,000 | 222 | 14 | 398 | 100 | 22,208 |
| Medium-70 | 1,345 | 768 | 1,999 | 212 | 8 | 398 | 100 | 21,184 |
| Medium-60 | 1,387 | 771 | 1,993 | 202 | 13 | 398 | 100 | 20,211 |
| Long-97 | 2,677 | 1,520 | 3,983 | 210 | 5 | 398 | 100 | 20,994 |
| Long-95 | 2,611 | 1,508 | 3,959 | 206 | 10 | 400 | 100 | 20,565 |
| Long-90 | 2,677 | 1,530 | 3,969 | 196 | 5 | 400 | 100 | 19,622 |
| Long-80 | 2,859 | 1,528 | 3,990 | 194 | 5 | 398 | 100 | 19,424 |
| Long-70 | 2,830 | 1,512 | 3,993 | 224 | 19 | 399 | 100 | 22,396 |
| Long-60 | 2,630 | 1,519 | 3,977 | 207 | 7 | 398 | 100 | 20,699 |
| Numerous-97 | 272 | 171 | 372 | 203 | 5 | 400 | 5,000 | 1,012,543 |
| Numerous-95 | 272 | 171 | 372 | 203 | 5 | 400 | 5,000 | 1,012,528 |
| Numerous-90 | 271 | 171 | 372 | 204 | 5 | 400 | 5,000 | 1,018,681 |
| Numerous-80 | 271 | 171 | 372 | 203 | 5 | 400 | 5,000 | 1,016,997 |

### Evaluation measures

A number of criteria for evaluating the results of clustering algorithms have been proposed (https://en.wikipedia.org/wiki/Clustemanalysis). Some criteria are applicable when the true clusters are known, other criteria are applicable when the true clusters are unknown, and others are applicable when the true clusters are available or unavailable.

#### Criteria for known true clusters

All (or the majority of) sequences in a good, predicted cluster should come from one real cluster. The purity criterion measures the mixing extent of sequences comprising a predicted cluster (Equation 6).

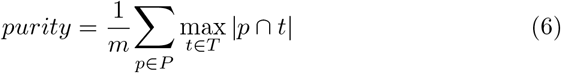

Here, *m* in the number of sequences comprising predicted clusters of size two or more, *P* is the set of predicted clusters, and *T* is the set of true clusters. The higher the purity, the better.

Jaccard index compares the number of sequences common to a predicted cluster and a real cluster (i.e., the intersection of the two clusters) to the total number of unique sequences found in both clusters (i.e., the union of the two clusters). Equation 7 describes how the Jaccard index is calculated. Perfect clustering results have a Jaccard score of 1.

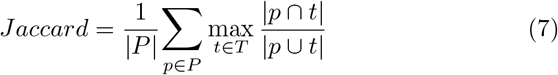

The G-Measure [35] is similar to the Jaccard index. It compares the number of common sequences to a predicted cluster and a real cluster to the size of each cluster separately (Equation 8). Perfect clustering results have a G-Measure score of 1.

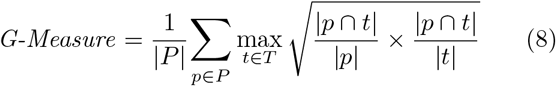

To look at these three criteria simultaneously, we define a new criterion called cluster quality for known clusters which is the harmonic mean of purity, Jaccard index, and G-Measure (Equation 9).

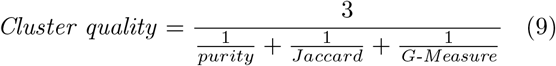

The harmonic mean is usually suitable to averaging rates that have the same range.

#### Criteria for unknown true clusters

In high-quality clustering, a cluster is desired to be compact and as far as possible from its neighbors to maximize separation. For each cluster, Davies-Bouldin index [36] evaluates the closest most-scattered cluster (Equation 10).

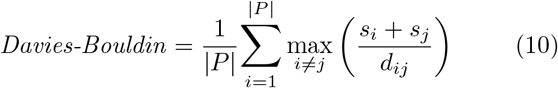

Here, *s_i_* measures the scatter of members of cluster *i* as the average distance between a member and the cluster center; *d_ij_* measures the separation between clusters i and *j* as the distance between their centers. The lower the Davies-Bouldin score, the better.

Dunn index [37] takes into account clusters’ scatter and separation. This index is calculated as the ratio of the distance between the closest two clusters in a data set to the maximum scatter (Equation 11). The higher the Dunn score, the better.

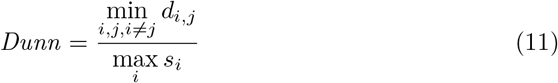

The Silhouette [38] criterion compares the placement of a sequence in a cluster to the placement of the same sequence in the closest neighbor cluster (12).

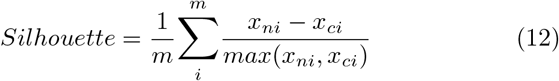

Here, *m* is the number of sequences in a data set; *x_ci_* is the distance between the *i^th^* sequence and the center of its own cluster and *x_ni_* is the distance between the *i^th^* sequence and the center of the closest neighboring cluster. Silhouette scores are between −1 (the worst score) and 1 (the best score). In our calculation of Silhouette scores, the neighbor cluster is determined for an entire cluster — not for each sequence separately — as the cluster with the closest center.

A distance is calculated as 1 minus the corresponding identity score reported by *Identity* [29]. Recall that an identity score is a sequence similarity measure calculated as the ratio of the number of identical nucleotides of two sequences to the length of the alignment (including gaps) of the two sequences.

Intra-cluster similarity is the average similarity (identity score) of a sequence to the center of its cluster (the higher, the better). Inter-cluster similarity is the average similarity between two centers of predicted clusters (the lower, the better).

To look at multiple criteria assessing the quality of predicted clusters, for which we do not have a ground truth, we defined the cluster quality for unknown clusters. The geometric mean is suitable to averaging values that have different unites or ranges. Thus, the cluster quality for unknown clusters is defined as the geometric mean of the following:

- 1 ÷ Bavies-Bouldin,
- Dunn,
- (1+ Silhouette) ÷ 2,
- Intra-cluster similarity, and
- 1– inter-cluster similarity.

#### Criteria for known and unknown true clusters

Coverage, run time, and memory requirement are criteria applicable to known and unknown true clusters. Coverage is the ratio of the total size of predicted clusters including at least two sequences to the size of the data set. The wall-clock time is reported. Memory is measured in Megabyte (MB).

## Results

### Results on synthetic data

#### Small data sets with 80–97% identity

We evaluated *CD-HIT, MeShClust v1.0, MeShClust v3.0*, and *UCLUST* on the following 12 testing data sets: Short 80, Short 90, Short 95, Short 97, Medium 80, Medium 90, Medium 95, Medium 97, Long 80, Long 90, Long 95, and Long 97. Each of these data sets includes less than 25,000 sequences. Note that these data sets were generated for this work and were not available at the time we developed *MeShClust v1.0*. Additionally, these *testing* data sets are different from the *training* data sets that were utilized while developing and optimizing *MeShClust v3.0*. We report the average performance on the 12 sets in Table 3 and the performance on each data set in Additional file 1.

**Table 3:** Evaluations on the synthetic testing data sets. The first set of evaluations was conducted on 12 sets, each of which includes clusters whose members are 80%, 90%, 95%, or 97% identical to a template sequence, i.e., a true center. The second set of evaluations was conducted on six data sets representing clusters of degenerate sequences (e.g., members are 60% or 70% identical to true centers). Each set of the first and second sets of evaluations includes less than 25k sequences. The third set of evaluations was conducted on four data sets, each of which includes more than one million sequences. Members comprising these clusters are 80%, 90%, 95%, or 97% identical to true centers. All clusters in the same data set have the same minimum identity score. For example, cluster members of the Short-97 data set are 97.00–99.99% identical to the true centers. The direction of the arrow below each criterion indicates whether a high or a low value is better.

| Tool | Purity<br>(↑) | Jaccard<br>(↑) | G-Measure<br>(↑) | Cluster quality<br>(↑) | Coverage<br>(↑) | Centers<br>(↑) | Time<br>(↓) | Memory (GB)<br>(↓) |
| --- | --- | --- | --- | --- | --- | --- | --- | --- |
| Short, Medium, and Long: 80–97% |  |  |  |  |  |  |  |  |
| *CD-HIT* | 0.92 | 0.19 | 0.33 | 0.29 | 0.92 | 0.01 | 00:71:00 | 0.36 |
| *MeShClust v1.0* | 0.99 | 0.92 | 0.93 | 0.94 | 0.99 | 0.35 | 00:00:26 | 0.20 |
| *MeShClust v3.0* | 1.00 | 1.00 | 1.00 | 1.00 | 1.00 | 0.78 | 00:05:18 | 6.55 |
| *MeShClust v3.0* (auto) | 1.00 | 1.00 | 1.00 | 1.00 | 1.00 | 0.80 | 00:12:08 | 6.59 |
| *UCLUST* | 0.70 | 0.08 | 0.20 | 0.15 | 0.70 | 0.00 | 00:00:16 | 0.12 |
| Short, Medium, and Long: 60–70% |  |  |  |  |  |  |  |  |
| *CD-HIT* | 1.00 | 0.14 | 0.28 | 0.24 | 0.90 | 0.01 | 01:23:46 | 0.35 |
| *MeShClust v1.0* | 0.98 | 0.93 | 0.96 | 0.96 | 1.00 | 0.53 | 00:00:25 | 0.19 |
| *MeShClust v3.0* | 1.00 | 1.00 | 1.00 | 1.00 | 1.00 | 0.76 | 00:11:44 | 5.79 |
| *MeShClust v3.0* (auto) | 1.00 | 1.00 | 1.00 | 1.00 | 0.98 | 0.65 | 00:14:32 | 5.83 |
| *UCLUST* | 1.00 | 0.22 | 0.34 | 0.34 | 0.83 | 0.01 | 00:00:28 | 0.08 |
| Numerous: 80–97% |  |  |  |  |  |  |  |  |
| *CD-HIT* | 1.00 | 0.48 | 0.59 | 0.62 | 1.00 | 0.00 | 00:39:31 | 0.91 |
| *MeShClust v1.0* | 1.00 | 0.81 | 0.83 | 0.87 | 0.99 | 0.01 | 00:19:04 | 2.58 |
| *MeShClust v3.0* | 1.00 | 0.99 | 0.99 | 1.00 | 1.00 | 0.15 | 02:58:15 | 12.76 |
| *MeShClust v3.0* (auto) | 1.00 | 0.98 | 0.99 | 0.99 | 0.99 | 0.15 | 02:50:40 | 13.06 |
| *UCLUST* | 1.00 | 0.07 | 0.20 | 0.15 | 0.89 | 0.00 | 00:06:41 | 0.72 |

We start by looking at cluster quality as defined in Equation 9, *MeShClust v3.0* came first, achieving almost perfect score (1.00); *MeShClust v1.0* came second (0.99) followed by *CD-HIT* (0.29) and *UCLUST* (0.15). The same trend was observed on the percentage of true centers identified by each tool (*MeShClust v3.0*: 78%; *MeShClust v1.0*: 35%; *CD-HIT*: 1%; and *UCLUST*: 0%). The greedy nature of *CD-HIT* and *UCLUST* does not allow them to identify the true centers. Because these data sets are synthetic, they do not include noisy sequences. Therefore, a coverage score of one is desired. A coverage score of less than one indicates that some sequences do not belong to any cluster. *MeShClust v3.0* and *MeShClust v1.0* achieved perfect or almost perfect coverage scores, followed by *CD-HIT* (0.92) and *UCLUST* (0.70). With respect to time, *UCLUST* was the fastest followed by *MeShClust v1.0, MeShClust v3.0*, and *CD-HIT. MeShClust v3.0* required the highest amount of memory. Regardless of the rank of *MeShClust v3.0* with respect to its time and memory requirements, these requirements are very reasonable. It took *MeShClust v3.0* five minutes and 18 seconds on average to process one of the 12 sets.

*MeShClust v3.0* required 6.55 GB of memory on average for each data set; such memory is available on almost all personal computers at the time of this writing. In sum, *MeShClust v3.0* achieved the highest scores on cluster quality and the percentage of true centers while requiring reasonable time and memory.

#### Small data sets with low identity

Clustering degenerate sequences (≤ 60% identity) using alignment-free statistics was a challenge while developing *MeShClust v1.0*. Therefore, we decided to utilize a global alignment algorithm in clustering degenerate sequences with *MeShClust v1.0*. Due to the improvement we introduced to *Identity* (the tool for calculating pairwise identity scores using alignment-free statistics), *MeShClust v3.0* is able to cluster degenerate sequences *without* the use of any alignment algorithms.

We evaluated the performances of the four tools on the following six testing data sets: Short 60, Short 70, Medium 60, Medium 70, Long 60, and Long 70. The average performance is reported in Table 3 and the performance on each data set in Additional file 1.

*MeShClust v3.0* achieved perfect score on the cluster-quality criterion; *MeShClust v1.0* came second with a score of 0.96, *UCLUST* came third with a score of 0.34, and *CD-HIT* came fourth with a score of 0.28. *MeShClust v3.0* and *MeShClust v1.0* were able to cluster almost all sequences, whereas *CD-HIT* and *UCLUST* clustered 90% and 83% of the sequences. *MeShClust v3.0* was able to identify 76% of the true centers, i.e., the predicted center of a cluster is the same as the true center, from which member sequences were generated. *MeShClust v1.0* identified 53% of the true centers, whereas *CD-HIT* and *UCLUST* identified 1%. With respect to processing time, *MeShClust v1.0* was the fastest followed by *UCLUST*, *MeShClust v3.0*, and *CD-HIT*. *MeShClust v3.0* took 11 minutes and 44 seconds on average to process one of the degenerate sets. The memory required by *MeShClust v3.0* was the highest (5.79 GB), whereas the three related tools required 0.08–0.35 GB. Overall, *MeShClust v3.0* is able produce high-quality clusters while utilizing available memory and acceptable processing time.

#### Large data sets with 80–97% identity

The previous two experiments were conducted on small data sets, each of which includes less than 25,000 sequences. Now, we report the performances of the four tools on the following testing sets: Numerous 80, Numerous 90, Numerous 95, and Numerous 97. Each of these data sets consists of more than one million sequences and 5,000 clusters. Table 3 displays the average performance on the four large sets. Additional file 1 includes the performance on each set individually.

According to the cluster-quality criterion, *MeShClust v3.0* achieved the highest score of 1.00; *MeShClust v1.0* came second with a score of 0.87; *CD-HIT* came third with a score of 0.62; *UCLUST* came fourth with a score of 0.15. According to the true-centers criterion, *MeShClust v3.0* identified the largest percentage of the true centers (15%), whereas *MeShClust v1.0* identified 1% and *CD-HIT* and *UCLUST* could not identify any true centers. Even though *MeShClust v3.0*’s performance based on the true-centers criterion is not as good as the performance on the small data sets, it is the best on the large data sets in contrast to the three related tools. According to the coverage criterion, *MeShClust v3.0* and *CD-HIT* achieved almost perfect coverage scores, followed *MeShClust v1.0* (0.99) and *UCLUST* (0.89). *MeShClust v3.0* took about three hours on average to process each of the large data sets. *MeShClust v3.0* was the slowest among the four tools. With respect to memory requirement, *MeShClust v3.0* required the largest amount of memory of 12.76 GB.

*MeShClust v3.0* achieved the highest scores on the cluster-quality, the true-centers, and the coverage criteria. Producing these high-quality clusters required more time and more memory than the related tools. From the prospective of a life scientist who is about to drive biological insights from sequence clusters, the high quality of clusters produced by *MeShClust v3.0* greatly outweighs its time and memory requirements, both of which are reasonable and available.

#### Estimating a threshold identity score from data

A life scientist may not know or uncertain of a good value (or at least a good starting point) for the threshold-score parameter. *CD-HIT*, *MeShClust v1.0*, and *UCLUST* require the user to provide a threshold identity score. *MeShClust v1.0* can handle some error in the threshold score; however, it still requires the user to provide a score. A tool uses this threshold as a cut-off for determining cluster membership. All members of a cluster must have identity scores with the center greater than or equal to the threshold score. *MeShClust v3.0* can estimate the threshold score without any assistance from the user.

First, we applied *MeShClust v3.0* to estimating threshold identity scores on the 22 *training* sets. The error ranged from 0.02% to 3.36% with an average of 1.05% and a standard deviation of 0.81%. Second, we applied *MeShClust v3.0* to estimating the threshold scores and using them in clustering the 22 *testing* sets. In Table 3, we report (marked with auto) the average performances in the three previous experiments. The performance due to an estimated threshold score is very comparable to that due to a user-provided one, demonstrating *MeShClust v3.0*’s success in estimating the threshold score needed for determining cluster membership. Another implication of these results is that a user may use *MeShClust v3.0* without adjusting any parameters, minimizing guesswork.

### Results on real data

We evaluated *CD-HIT*, *MeShClust v1.0*, *MeShClust v3.0*, and *UCLUST* on the Microbiome data set, which includes more than one million sequences, using an identity score of 97% (Table 4). *MeShClust v3.0* outperformed all related tools on every criterion measuring cluster quality. According to the overall cluster quality measure, *MeShClust v3.0* came first followed by *MeShClust v1.0, CD-HIT*, and *UCLUST. MeShClust v3.0* achieved 100% improvement over our previous version, more than 200% over *CD-HIT*, and 300% over *UCLUST*. The coverage of our tool was 96%, meaning that 4% of the sequences did not belong to any clusters according to the 97% threshold. One possible explanation is that the unclustered sequences represent noisy sequences due to sequencing errors. The improvement in *MeShClust v3.0*’s cluster quality came at the cost of time and memory. *MeShClust v3.0* took the longest time (one hour and 41 minutes) and required the largest memory (about 15 GB), whereas the other related tools took few minutes and required 0.42–2.56 GB of memory. *MeShClust v3.0*’s memory requirement of 15 GB is already available on almost all personal computers at the time of this writing. When scientific conclusions depend on the formed clusters, waiting for two hours to obtain high-quality clusters is well justified.

**Table 4:** Evaluations on real data sets. The microbiome set was clustered with identity score of 97%. The LTRs set was clustered with identity score of 70%. The direction of the arrow below each criterion indicates whether a high or a low value is better.

| Tool | Dunn<br>(↑) | Davies-Bouldin<br>(↓) | Silhouette<br>(↑) | Intra<br>(↑) | Inter<br>(↓) | Cluster quality<br>(↑) | Coverage<br>(↑) | Time<br>(↓) | Memory (GB)<br>(↓) |
| --- | --- | --- | --- | --- | --- | --- | --- | --- | --- |
| Microbiome |  |  |  |  |  |  |  |  |  |
| *CD-HIT* | 0.01 | 3.39 | 0.24 | 0.94 | 0.94 | 0.16 | 0.99 | 00:01:45 | 0.93 |
| *MeShClust v1.0* | 0.02 | 1.50 | 0.43 | 0.96 | 0.90 | 0.25 | 0.99 | 00:05:01 | 2.56 |
| *MeShClust v3.0* | 0.27 | 0.77 | 0.74 | 0.97 | 0.90 | 0.50 | 0.96 | 01:41:06 | 15.08 |
| *UCLUST* | 0.01 | 5.50 | -0.14 | 0.94 | 0.96 | 0.12 | 0.98 | 00:00:31 | 0.42 |
| LTRs |  |  |  |  |  |  |  |  |  |
| *CD-HIT* | 0.02 | 1.81 | 0.05 | 0.78 | 0.52 | 0.29 | 1.00 | 3:47:05 | 0.75 |
| *MeShClust v1.0* | 0.14 | 1.13 | 0.33 | 0.86 | 0.52 | 0.51 | 1.00 | 2:57:07 | 1.75 |
| *MeShClust v3.0* | 0.98 | 0.86 | 0.47 | 0.88 | 0.58 | 0.79 | 0.94 | 2:01:22 | 16.24 |
| *UCLUST* | 0.02 | 2.02 | 0.11 | 0.75 | 0.55 | 0.27 | 1.00 | 9:01:14 | 0.50 |

Next, we evaluated the tools on the LTRs data set, which includes about 250,000 sequences, using an identity score of 70%. This data set was produced by a program called LtrDetector that discovers LTR retrotransposons [39]. LtrDetector processed the maize genome. One way to reduce the false positive rate of LtrDetector is to cluster the LTRs; sequences with predicted LTRs that do not belong to any cluster are likely to be false positives and should be removed. *MeShClust v3.0* outperformed the three related tools on every criterion related to cluster quality except the inter-cluster similarity measure. According to the overall cluster quality measure, our tool came first followed by *MeShClust v1.0, CD-HIT*, and *UCLUST*. The margin of improvement is very clear (55% over *MeShClust v1.0*, 172% over *CD-HIT*, and 193% over *UCLUST*). Only 94% of the LTRs were clustered. This coverage is in line with the assumption that this data set includes false positives; the purpose of clustering these sequences is to identify and remove sequences that do not belong to any cluster. Similar to the Microbiome data set, our tool took the longest time and required the largest amount of memory. However, our time and memory requirements are well justified given the high quality of the clusters produced by *MeShClust v3.0*.

In the third evaluation experiment on real data, we evaluated the four tools on the viral data set with a threshold identity score of 50%. This is a challenging experiment because k-mer statistics may be ineffective on highly degenerate sequences [40]. To circumvent this challenge while developing *MeShClust v1.0*, we utilized the actual global alignment algorithm instead of the k-mer-statistics-based classifier when the threshold identity score is 60% or below. *MeShClust v3.0* is completely free from alignment algorithms and depends entirely on alignment-free k-mer statistics. The results of this experiment are shown in Table 5. Recall that the viral data set consists of nine known clusters. In terms of cluster quality, *MeShClust v1.0* (0.81) came first followed by *CD-HIT* (0.78), *MeShClust v3.0* (0.71), and *UCLUST* (0.43). In terms of coverage, the same exact performance ranking was observed. In terms of time, *CD-HIT* was the fastest followed by *MeShClust v3.0*, *UCLUST*, and *MeShClust v1.0*. In terms of memory, *MeShClust v1.0* and *UCLUST* came first, *MeShClust v3.0* came second, and *CD-HIT* came last. Although *MeShClust v3.0* did not come first in this experiment, its cluster quality is within 10–14% from those obtained by the first and the second-best performing tools. Because clustering degenerate sequences using alignment-free k-mer statistics could not be done in the past, these results represent a progress in the alignment-free field.

In the fourth evaluation experiment on real data, we applied *MeShClust v3.0* to clustering bacterial genomes. *None of the related tools can cluster very long sequences such as those of bacteria because of the prohibitive time alignment algorithms would take to calculate pairwise identity scores*. We applied our tool to clustering bacterial genomes into species. To start, we assembled the 14-bacterial species data set, which includes 1,328 sequences representing 14 bacterial species. The length of an average genome is approximately 3 mega base pairs. Because this is the first time to cluster bacterial genomes using identity scores, there is no community-accepted threshold identity score as the one we utilized in clustering the microbiome data set (97%). Using the threshold-estimation feature of *MeShClust v3.0*, an identity score of 91.23% served as a starting point. The clusters obtained due to the estimated threshold were fine but not perfect — their cluster quality was 0.77 (a perfect cluster quality score is 1). After that, we tried three lower threshold scores of 90%, 85%, and 80%. Table 5 shows the results of this experiment. The clusters due to the 80% and 85% threshold scores had high cluster quality and high coverage, suggesting that bacterial genomes of the same species are 80-85% identical to each other. Using a threshold identity score of 84%, we clustered the bacterial data set, which consists of 10,562 sequences with an average length of 3.7 mega base pairs. This experiment represents a stress test for *MeShClust v3.0* because of the high time and memory requirements. *MeShClust v3.0* successfully clustered this data set in about 50 hours using 56 GB of memory, resulting in 482 clusters of size 2 or greater covering 67% of the data set. To the best of our knowledge, this is the first time long sequences such as those of bacterial genomes can be clustered using identity scores.

**Table 5:** Evaluations on the viral and the 14-bacterial-species data sets. The viral set was clustered with identity score of 50%; it includes nine clusters representing nine viruses. The 14-bacterial-species set was clustered with multiple identity scores; it includes 14 clusters representing 14 bacterial species.

| Tool | Purity<br>(↑) | Jaccard<br>(↑) | G-Measure<br>(↑) | Cluster quality<br>(↑) | Coverage<br>(↑) | Time<br>(↓) | Memory (GB)<br>(↓) |
| --- | --- | --- | --- | --- | --- | --- | --- |
| Viral data set |  |  |  |  |  |  |  |
| *CD-HIT* | 0.97 | 0.67 | 0.77 | 0.78 | 0.95 | 00:00:02 | 0.18 |
| *MeShClust v1.0* | 0.91 | 0.72 | 0.83 | 0.81 | 0.98 | 00:00:28 | 0.08 |
| *MeShClust v3.0* | 0.96 | 0.56 | 0.72 | 0.71 | 0.72 | 00:00:07 | 0.12 |
| *UCLUST* | 1.00 | 0.26 | 0.46 | 0.43 | 0.64 | 00:00:17 | 0.08 |
| 14-bacterial-species set |  |  |  |  |  |  |  |
| *MeShClust v3.0* (0.80) | 1.00 | 1.00 | 1.00 | 1.00 | 1.00 | 00:29:36 | 14.09 |
| *MeShClust v3.0* (0.85) | 1.00 | 0.93 | 0.97 | 0.97 | 0.91 | 00:46:01 | 14.11 |
| *MeShClust v3.0* (0.90) | 1.00 | 0.64 | 0.76 | 0.78 | 0.96 | 00:42:54 | 14.21 |
| *MeShClust v3.0* (auto) | 1.00 | 0.62 | 0.73 | 0.77 | 0.93 | 02:48:41 | 14.21 |

## Discussion

### The scaled-up mean shift algorithm in action

Figure 2 shows detail about how *MeShClust v3.0* discovered centers in the Numerous-97 training data set, which includes 5,000 clusters and more than one million sequences. The bottom plot of Figure 2 shows the number of sequences read as *MeShClust v3.0* progresses. Recall that *MeShClust v3.0* employs an out-of-core strategy, in which the tool processes input sequences batch by batch. *MeShClust v3.0* scanned the data three times until it converged. The middle figure shows the number of sequences that do not belong to any of the clusters discovered so far; these sequences are kept in the reservoir. The reservoir is utilized during the first data pass only; therefore, it is empty during the second and third data passes. The top plot shows the number of discovered centers as the tool runs. All centers are discovered in the first data pass; the first set of centers were discovered by the main mean shift instance that was started with a quadratic initialization step. Additional centers were discovered by running an independent instance of the mean shift with a quadratic initialization step when 25,000 (the batch size) sequences were accumulated in the reservoir. For this reason, every increase in the number of centers is associated with a decrease in the reservoir size due to removing a batch. The number of discovered centers decreased a little bit (from 5,016 to 5,012) during the second data pass because some centers were merged. The number of centers did not change at all during the third data pass, resulting in convergence.

**Figure 2:**
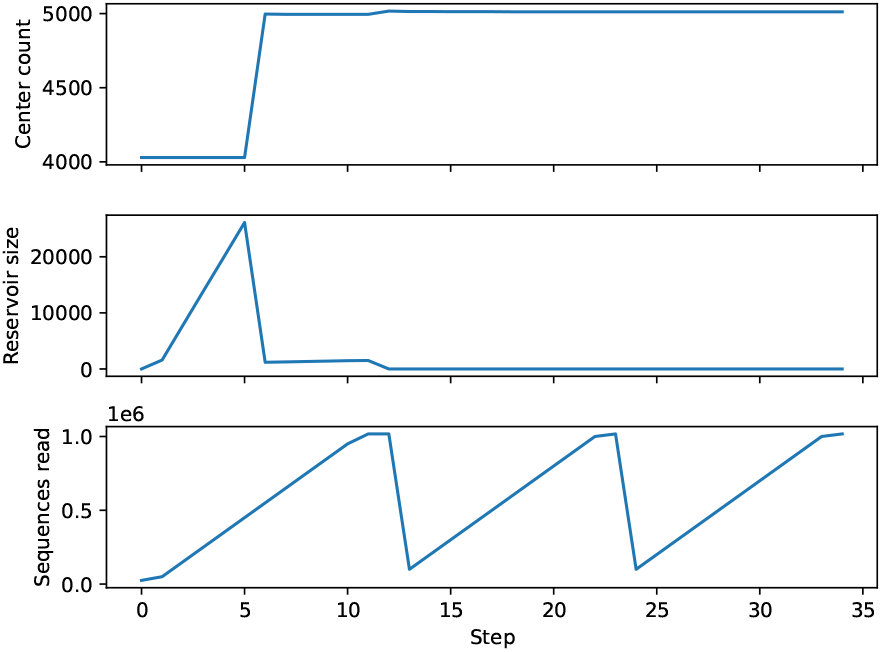
*MeShClust v3.0* in action on the Numerous-97 training data set. The top plot shows the number of centers as the algorithm runs. The middle plot shows the number of sequences accumulated in the reservoir; this number changes in the first data pass and is zero in the second and third data passes. The bottom plot shows the number of sequences read during the three data passes.

### Effects of the all-vs-all block size on *MeShClust v3.0*

To evaluate the effects of the all-vs-all block size on the performance of *MeShClust v3.0*, we ran our tool using different block sizes (1k, 2k, 5k, 10k, 15k, 20k, 25k, and 46k) and measured cluster quality, percentage of true centers, coverage, time, and memory (Figure 3). Recall that this block of sequences is used for initializing the mean shift algorithm, which applies *Identity* to calculating the similarity scores of every unique pair of sequences in this block. In other words, *Identity* performs all-vs-all on this block of sequences. This step is the most time-consuming step in *MeShClust v3.0*. The larger the block size is, the more time and memory are required. We performed this experiment on four training data sets: 60-Short, 70-Medium, 80-Long, and 97-Numerous. Each of the first three data sets includes less than 25k sequences, whereas the 97-Numerous set includes more than one million sequences.

**Figure 3:**
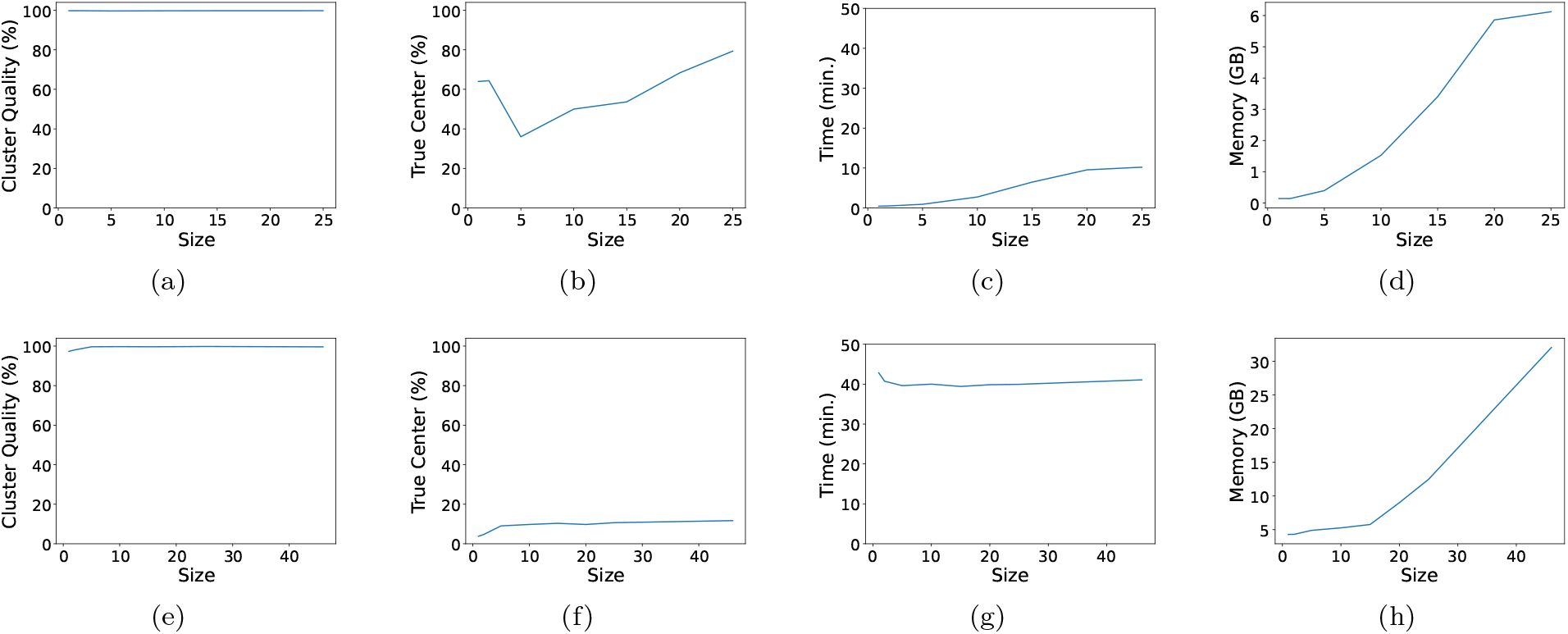
The effects of the size of the all-vs-all block on cluster quality, percentage of true centers, time, and memory. Figures a–d are produced by evaluating *MeShClust v3.0* using different block sizes (1k, 2k, 5k, 10k, 15k, 20k, and 25k) on three small data sets: 60-Short, 70-Medium, and 80-Long; each of these sets consists of less than 25k sequences and includes 100 clusters. Figures e–h are produced by evaluating *MeShClust v3.0* using different block sizes (1k, 2k, 5k, 10k, 15k, 20k, 25k, and 46k) on one large data set (the 97-Numerous set), which includes more than one million sequences and 5,000 clusters.

First, we looked at cluster quality (the harmonic mean of purity, Jaccard index, and G-Measure). On the small data sets (60-Short, 70-Medium, and 80-Long), the block size seems to have no effect on cluster quality. On the large data set (97-Numerous), we observed a minor increase in performance as the block size increased; almost the same cluster qualities were obtained using a size of 10k or larger. After that we looked at the percentage of true centers identified. On the small data sets, using large blocks (20–25k) resulted in identifying more true centers than using small sizes. Same trend was observed on the large data set, but it was not as strong as that observed on the small data sets. The block size has almost no effect on the coverage measure (99.6–99.7% on average on the three small data sets and 99.6–99.9% on the large data set).

With respect to time, *MeShClust v3.0* took longer time using large blocks than using small sizes on the three small data sets. This time trend is expected because *MeShClust v3.0* utilized one mean shift instance with the quadratic initialization step while processing the small sets. On the large data set, the time requirements due to different block sizes were very similar. Processing small blocks takes shorter time and discovers a smaller number of new centers than processing large blocks. The smaller the number of discovered centers is, the more frequent executing the mean shift with the quadratic initialization step occurs and vice versa, i.e., these two trends balance each other. Finally, we looked at memory, which increases as the block size increases.

In sum, using a large block size is likely to result in identifying more true centers than using a small size but at the cost of memory. Further, cluster quality due to a large block size is likely to be better than that due to a small block size — specially on large data sets.

### *MeShClust v1.0* versus *MeShClust v3.0*

Even though *MeShClust v1.0* and *MeShClust v3.0* are both based on the mean shift algorithm, their adaptations of the algorithm are completely different. *MeShClust v1.0* consists of two stages. In the first stage, it builds initial clusters one at a time. First, *MeShClust v1.0* sorts its input sequences according to length. Starting with the shortest sequence as the initial center of the first cluster, it scans all input sequences for sequences that are similar to the center. The mean of these sequences is calculated (sequences are converted into k-mer histograms) and a representative sequence is selected. This representative sequence becomes the new center of the cluster. The just-found, similar sequences are removed from the input sequences. Next, *MeShClust v1.0* scans the remaining sequences for sequences that are similar to the new center. This procedure — (i) find similar sequences, (ii) calculate a new mean, and (iii) find a representative sequence — is repeated until no new similar sequences can be found. At that time, the cluster is put aside, and a new sequence is selected from the remaining sequences to be the initial center of the new cluster. The new sequence is the closest sequence (among the remaining sequences) to the center of the previous cluster. The first stage results in a semi-sorted list of clusters. In the second stage, *MeShClust v1.0* attempts to merge a cluster with the five clusters before it in the list and the five clusters after it in the list. In contrast to *MeShClust v1.0, MeShClust v3.0* builds many clusters at the same time — not only one. *MeShClust v3.0* attempts to merge a cluster with every other cluster — not only the neighboring 10 clusters. *MeShClust v3.0* may scan input sequences as many times as needed until the algorithm converges — not only two stages. *MeShClust v3.0* processes input data batch by batch — the entire data set is *not* loaded in memory. For this reason, *MeShClust v1.0* cannot process very large data sets.

*MeShClust v3.0* is alignment free, whereas *MeShClust v1.0* is not entirely alignment free. *MeShClust v1.0* determines sequence similarity using a machinelearning-based classifier that is trained on identity scores produced by a global alignment algorithm. *MeShClust v3.0* utilizes *Identity* whose core is a regression model trained on identity scores that are calculated by introducing mutations into real sequences, i.e., alignment algorithms are never used. For this reason, *MeShClust v3.0* can be applied to very long sequences, e.g., bacterial genomes, but *MeShClust v1.0* cannot be applied because of the impractical long time a global alignment algorithm would take on such long sequences. When the threshold identity score is less than or equal to 60%, *MeShClust v1.0* uses a global alignment algorithm — instead of the classifier — to determine the similarity between a sequence and the center of a cluster, whereas *MeShClust v3.0* uses *Identity* which is completely alignment free.

The classifier utilized in *MeShClust v1.0* depends on four features only and applies a greedy strategy for feature selection, whereas *MeShClust v3.0* utilizes *Identity* that employs the best-first algorithm to select features from 903 statistics, i.e., a stronger feature-selection algorithm and many informative features.

### MeShClust v2.0

The adaptation of the mean shift algorithm in *MeShClust v2.0* [41] is the same as that of *MeShClust v1.0*. The difference between the two versions is that the second version’s classifier utilized in selecting similar sequences does not utilize alignment algorithms to generate identity scores for training; it uses similar idea to that implemented in *Identity*.

## Conclusions

Computational tools for clustering DNA sequences are utilized in many studies in molecular biology. Two pioneering tools — *CD-HIT* and *UCLUST* — are based on greedy algorithms. For this reason, they may result in fragmented clusters and are unable to identify the true centers of the clusters they form. Further, these widely used tools cannot be applied to clustering long sequences such as those of bacterial genomes. Earlier, we developed *MeShClust v1.0*, which is based on the mean shift algorithm to address some of the limitations of *CD-HIT* and *UCLUST*. Although the adaptation of the mean shift algorithm in *MeShClust v1.0* was a step forward, it was not the original algorithm.

We are convinced that applying the original mean shift algorithm should result in high-quality clusters. In this work, we propose *MeShClust v3.0*, which applies the original mean shift algorithm on small data sets and a scaled-up version of it on large data sets.

The main contributions of *MeShClust v3.0* are:

- High-quality clusters;
- Clustering long sequences as those of bacterial genomes using identity scores for the first time;
- Clustering large data sets that cannot fit in memory using the mean shift algorithm;
- Automatic estimation of the threshold identity score; and
- Progress towards using alignment-free k-mer statistics in clustering DNA sequences.

*MeShClust v3.0* represents progress in terms of cluster quality and scale, resulting in accurate biological insights and providing opportunities for new studies.

## Availability and requirements

**Project name**: *MeShClust v3.0*

**Project home page**: https://github.com/BioinformaticsToolsmith/Identity

**Operating system(s)**: UNIX/Linux/Mac **Programming language**: C++

**Other requirements**: GNU g++ 7.5.0 or later, GNU Make, and CMake v3.10

**License**: Affero General Public License version 1

**Any restrictions to use by non-academics**: Alternative commercial license is required

## List of abbreviations

LTR: Long Terminal Repeat.

## Additional Files

Additional file 1 — Results on the synthetic testing data sets

This file includes the results of evaluating *CD-HIT, MeShClust v1.0, MeShClust v3.0*, and *UCLUST* on all synthetic testing data sets. Results shown in Table 3 were derived from this file. File format is MS Excel (.xlsx).

